# Determining the structure of protein-bound ceramides, essential lipids for skin barrier function

**DOI:** 10.1101/2023.08.03.551778

**Authors:** Yusuke Ohno, Tetsuya Nakamura, Takafumi Iwasaki, Akira Katsuyama, Satoshi Ichikawa, Akio Kihara

## Abstract

Protein-bound ceramides, specialized ceramides covalently bound to corneocyte surface proteins, are essential for skin permeability barrier function. However, their exact structure and target amino acid residues are unknown. Here, we found that epoxy-enone (EE) ceramides, precursors of protein-bound ceramides, as well as their synthetic analog, formed stable conjugates only with Cys among nucleophilic amino acids. NMR spectroscopy revealed that the β-carbon of the enone was attached by the thiol group of Cys via a Michael addition reaction. We confirmed the presence of Cys-bound EE ceramides in mouse epidermis by mass spectrometry analysis of protease-digested epidermis samples. EE-ceramides were reversibly released from protein-bound ceramides via sulfoxide elimination. We found that protein-bound ceramides with reversible release properties accounted for approximately 60% of total protein-bound ceramides, indicating that Cys-bound EE ceramides are the predominant protein-bound ceramides. Our findings provide clues to the molecular mechanism of skin barrier formation by protein-bound ceramides.

**Highlights:**

- Epoxy-enone (EE) ceramides form stable conjugates with Cys.
- Cys binds to the β-carbon of the enone via a Michael addition reaction.
- Cys-bound EE ceramides exist in the mouse epidermis.
- Cys-bound EE ceramides are the predominant protein-bound ceramides.

**In Brief:** Protein-bound ceramides are specialized ceramides essential for skin permeability barrier function. In the present study, we determined the exact structure of protein-bound ceramides as Cys-bound epoxy-enone ceramides where the β-carbon of the epoxy-enone ceramides is attached by Cys residues of corneocyte surface proteins via a Michael addition reaction.

## Introduction

The skin permeability barrier (skin barrier) protects our bodies against invasion of external material, including pathogens, allergens, and chemicals, and prevents the loss of internal water and electrolytes. Abnormalities in skin barrier function increase the risk of or cause infectious diseases, atopic dermatitis, and ichthyosis, a skin disorder characterized by dry, thickened, and scaly skin.^1-4^ The stratum corneum, which constitutes the outermost layer of the epidermis, plays a pivotal role in skin barrier function and contains two key lipid structures: a lipid lamella and a corneocyte lipid envelope (CLE). The lipid lamella is a multilayered lipid structure that fills the space between corneocytes and is composed of ceramides, cholesterol, and free fatty acids (FAs).^5,6^ CLE is a membrane structure covering corneocytes; its constituents are specialized ceramides, called protein-bound ceramides.^7-9^

Ceramides are composed of a long-chain base (LCB) and an amide-linked FA. Numerous ceramide classes with different combinations of LCBs and FAs are found in mammalian epidermis.^10,11^ Ceramides are categorized into free ceramides (non-protein-bound ceramides) and protein-bound ceramides; free ceramides are further classified into non-acylated ceramides (conventional ceramides existing in a wide range of tissues) and acylceramides (ω-*O*-acylceramides). The FA moiety of acylceramides is hydroxylated, and linoleic acid is attached to the ω-hydroxyl group via an ester linkage. This unique acylceramide structure with three hydrophobic chains forms and maintains lipid lamellae.^12,13^ Protein-bound ceramides are covalently bound to proteins of the cornified envelope (CE), a cross-linked protein structure.^7-9,14^ Protein-bound ceramides have functions to confer resistance to chemicals and surfactants to corneocytes and stably connect corneocytes and lipid lamellae. Both acylceramides and protein-bound ceramides are crucial for skin barrier function. Mutations in the genes involved in their synthesis cause congenital ichthyosis in humans,^15-24^ and the respective gene knockout (KO) mice exhibit neonatal lethality due to skin barrier defects.^12,13,25-32^

Two models have been proposed for the structure of protein-bound ceramides. In the first model, the ω-hydroxyl group of ceramides with ω-hydroxy (ω-OH) FA (ω-OH ceramides) is ester-linked to the Glu residues of CE proteins (ceramides with protein-bound ω-OH FA [P-O ceramide] model).^33^ In the second model, the linoleic acid moiety of the acylceramides is modified to epoxy-enone, then the enone is covalently bound to the CE proteins by a Michael addition reaction or Schiff base formation (ceramides with protein-bound FA-esterified ω-OH FA [P-EO ceramide] model).^31^ In the experiment that provided the basis for the P-O ceramide model, the protein-bound ceramide fraction (the remnant after extraction of free lipids with organic solvents) was subjected to mild saponification, proteinase K treatment, recovery of peptide-bound ceramides by C4 column, cleavage of the bonds between peptides and ceramides by complete saponification, and mass spectrometry (MS) analyses.^33^ However, the amounts of protein-bound ceramides (P-O ceramides) detected by this analysis only accounted for 15–20% of total protein-bound ceramides, leaving the structure/binding mode of the remaining protein-bound ceramides unresolved.

P-O ceramides are considered to be produced via the following pathway. First, the linoleic acid moiety of acylceramides undergoes peroxidation and subsequent epoxy-hydroxylation by successive reactions catalyzed by the lipoxygenases ALOX12B and ALOXE3, respectively, producing epoxy-hydroxy ceramides (Figure 1).^34^ Both *ALOX12B* and *ALOXE3* are the causative genes of congenital ichthyosis.^16^ Epoxy-hydroxy ceramides are then cleaved by unknown lipases, either directly or after being further modified to triol ceramides or epoxy-enone (EE) ceramides, releasing ω-OH ceramides. Finally, the ω-OH group of the ω-OH ceramides are attached to CE proteins, generating P-O ceramides.^35^

**Figure 1.**
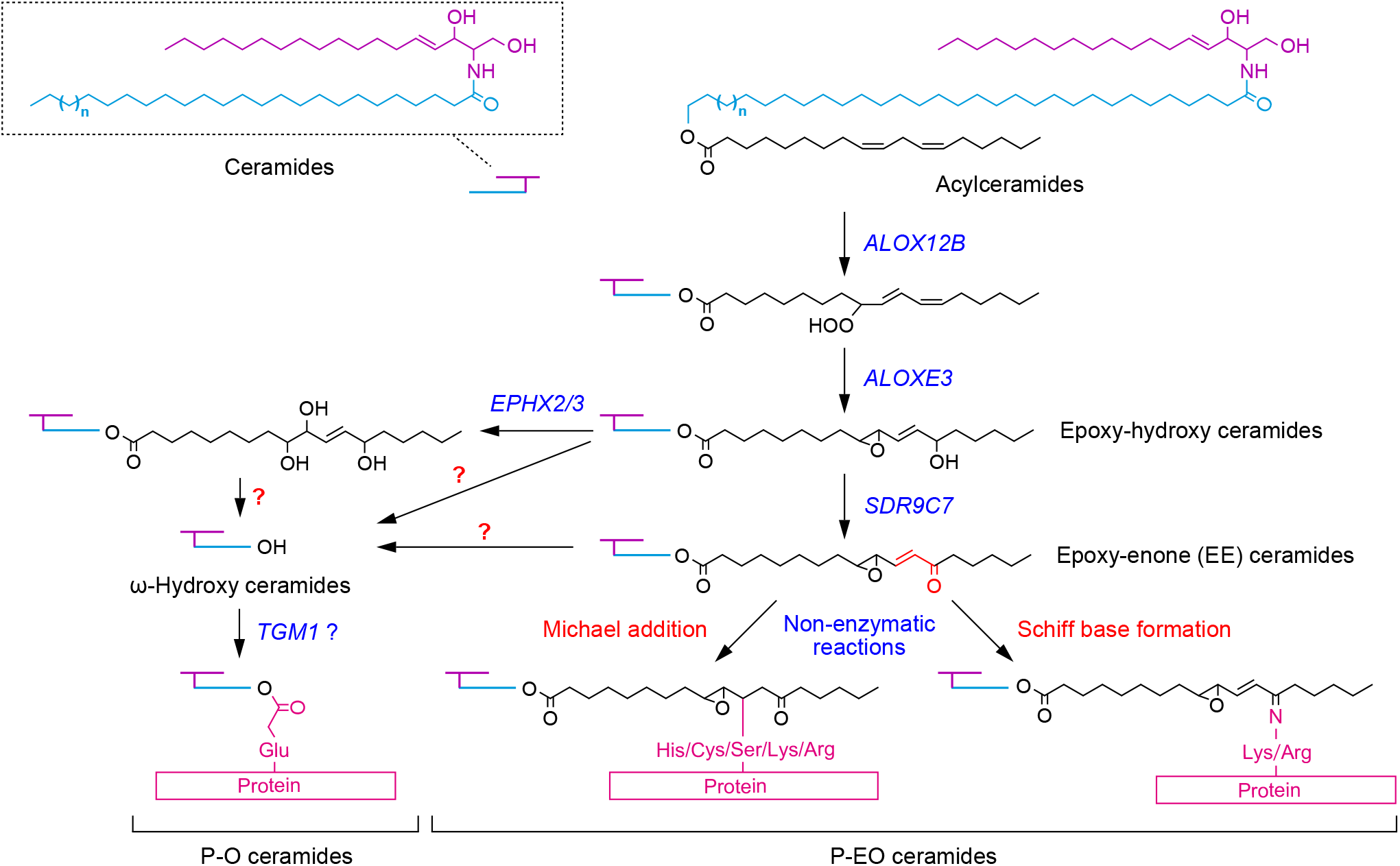
Two structural models and synthetic pathway of protein-bound ceramides. Two structural models of protein-bound ceramides (P-O ceramides and P-EO ceramides), their synthetic pathway, and genes involved in each reaction step are shown. A portion of acylceramides is converted to protein-bound ceramides after modification of their linoleic acid moiety. The linoleic acid moiety undergoes peroxidation, then epoxy-hydroxylation, producing epoxy-hydroxy ceramides. In the P-O ceramide synthetic pathway, epoxy-hydroxy ceramides are cleaved by unknown lipases, either directly or after being further modified to triol ceramides or EE ceramides, releasing ω-OH ceramides. Subsequently, the ω-OH group of ω-OH ceramides is attached to CE proteins, generating P-O ceramides. In the P-EO ceramide synthetic pathway, epoxy-hydroxy ceramides are reduced to EE ceramides. The α,β-unsaturated carbonyl (enone) group of EE ceramides then binds to the nucleophilic amino acids by a Michael addition reaction or a Schiff base formation, presumably in a non-enzymatic manner.

The P-EO ceramide model was recently proposed based on the finding that the reductase encoded by the ichthyosis causative gene *SDR9C7* converts epoxy-hydroxy ceramides to EE ceramides.^31^ Enones (α,β-unsaturated carbonyl groups) are highly reactive in general and can react and bind with proteins with Lys and Arg residues via a Schiff base formation, and with other nucleophilic amino acid residues (Cys, Ser, and His) via a Michael addition reaction.^36-38^ Therefore, P-EO ceramides are presumed to be generated by the conjugation of the enone moiety of EE ceramides with CE proteins via a Schiff base formation or a Michael addition reaction. However, the existence of P-EO ceramides in the epidermis has not yet been proven experimentally, and their binding mode (Schiff base formation and/or Michael addition reaction), as well as the target amino acids/proteins of EE ceramides, remain elusive.

To date, no methods for detecting/quantifying P-O ceramides/P-EO ceramides in the peptide- or amino acid-bound state have yet been established due to technical difficulties. Instead, ω-OH ceramides released from P-O/P-EO ceramides by hydrolyzing the ester bond with alkali are measured. However, this method cannot discriminate between P-O ceramides and P-EO ceramides, leaving the presence and quantity of each unclear.

In this study, we established a detection/quantification method for P-EO ceramides and revealed the target amino acid residue (Cys) and reaction mode (Michael addition reaction) of EE ceramides. Furthermore, we show that Cys-bound P-EO ceramides are the predominant protein-bound ceramides. These findings provide an important basis for elucidating the molecular mechanisms of skin barrier formation and for developing diagnosis and therapy for skin diseases caused by abnormal skin barrier formation.

## Results

### Binding of EE-analog with Cys

To examine whether EE ceramides bind to nucleophilic amino acids, we first used a chemically-synthesized analog of EE ceramides (EE-analog; Figure 2A). After incubating the EE-analog with Cys, Ser, His, Arg, and Lys, the reaction products were separated by thin-layer chromatography (TLC), followed by copper phosphate-staining. Only when incubated with Cys, the EE-analog band disappeared, concomitant with the appearance of a new band with lower mobility than that of the EE-analog band (Figure 2B). To gain some understanding of the structure of this new band, we performed ninhydrin staining, which marks primary amines as purple and secondary amines as yellow. The reaction product of EE-analog with Cys was detected in purple (Figure 2C), indicating the presence of a primary amine. Therefore, Cys was bound to the EE-analog via a thiol group, not an amino group.

**Figure 2.**
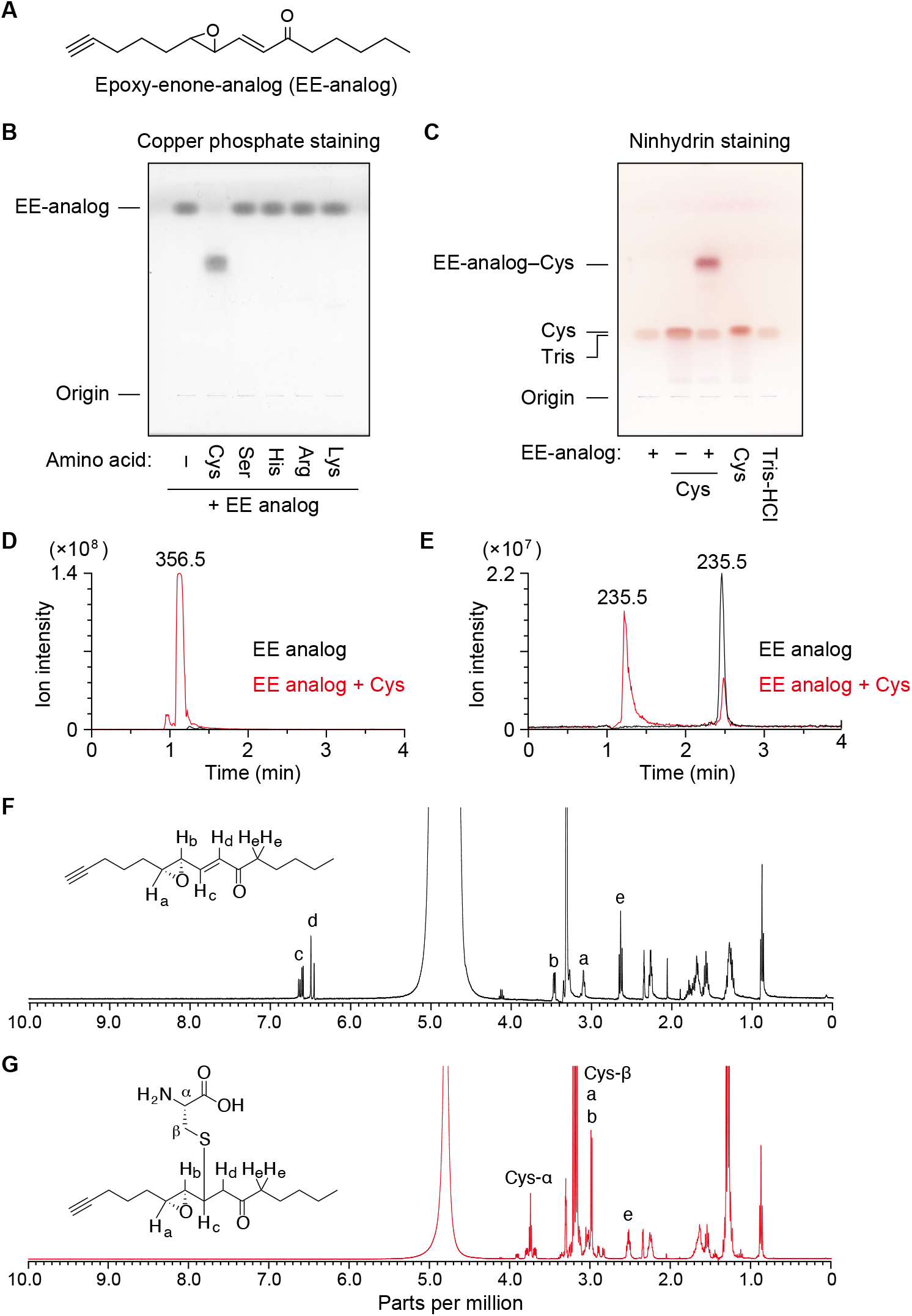
Conjugation of EE-analog and Cys. (A) Structure of the synthetic analog of EE ceramides (EE-analog). (B and C) EE-analog (2 mM, dissolved in CH_3_OH) was mixed with an amino acid (Cys, Ser, His, Arg, or Lys; 2 mM each, dissolved in 50 mM Tris-HCl [pH 7.4]) and incubated at 37 °C for 1 h. The reaction products (B and C), EE-analog (B and C), Cys (C), and reaction buffer (Tris-HCl; C) were separated by TLC and visualized by copper phosphate staining (B) or ninhydrin staining (C). (D and E) 2 mM EE-analog was mixed with 2 mM Cys and incubated at 37 °C for 1 h. The molecular ions with *m/z* 356.5 (D) and 235.5 (E) were detected via LC-MS using a single ion monitoring mode. (F and G) EE-analog only (44 mM, dissolved in CD_3_OD; F) or mixture of EE-analog and Cys (112 mM, dissolved in D_2_O containing 112 mM triethylamine; G) were incubated at 25 °C for 5 min and subjected to ^1^H NMR spectroscopy (400 MHz). The signals of the chemical shifts corresponding to the protons in each structure are indicated by letters in NMR spectra. See also Figure S5.

To confirm the reaction between the EE-analog and Cys, we conducted liquid chromatography (LC) coupled mass spectrometry (MS) by selecting molecular ions with *m/z* values of 356 and 235 corresponding to the conjugate of EE-analog and Cys (EE-analog–Cys) and the proton adduct of EE-analog, respectively. We detected a molecular ion peak of EE-analog-Cys when the EE-analog was incubated with Cys (Figure 2D). Moreover, this peak was not detected by EE-analog alone. The intensity of the EE-analog molecular ion peak (LC retention time, 2.5 min) was greatly reduced by incubation with Cys (Figure 2E). Another peak was also detected at an earlier retention time (1.2 min) than that of the EE-analog peak when the EE-analog was incubated with Cys. Since this peak had the same retention time as EE-analog–Cys (Figure 2D), it should represent the in-source decay product of EE-analog–Cys, i.e., EE-analog–Cys being converted to EE-analog during ionization.

To determine the structure of EE-analog–Cys, ^1^H NMR analysis was performed. The signals corresponding to the olefinic protons of an α,β-unsaturated ketone (c and d; Figure 2F) disappeared following incubation of EE-analog with Cys for 5 min, indicating that the nucleophilic addition occurred rapidly at the β-carbon. In addition, disappearance of the conjugated olefin led to an upfield shift of signal b. We excluded the possibility of a ring opening of the epoxide or 1,2-addition to the ketone, as the chemical shifts of protons a and e, which are adjacent protons of the reaction center for each reaction, barely changed following the EE-analog reaction with Cys. From these results, we concluded that EE-analog forms a stable adduct with only Cys among nucleophilic amino acids and that the adduct is formed by a Michael addition of the thiol group of Cys to the β-carbon of the enone of the EE-analog.

### Effect of functional groups in EE-analog on the formation of EE-analog–Cys conjugate

To examine the requirements of keto and epoxy groups in EE-analog binding to Cys, we prepared an alcohol form of EE-analog (epoxy-enol analog) in which the keto group of EE-analog was reduced; a dihydroxy form (dihydroxy-enone analog) with a cleaved epoxy group; and an α,β-unsaturated carbonyl compound (2-octen-4-one) without an epoxy group (Figure 3A). Except for commercially available 2-octen-4-one, epoxy-enol and dihydroxy-enone analogs were synthesized from the EE-analog by chemical (reduction with NaBH_4_) and enzymatic (cleavage with epoxide hydrolase EPHX2) reactions, respectively. TLC analysis confirmed high purity of the synthesized compounds (Figure S1A). Epoxy-enol analog was detected as two bands in close proximity on the TLC plate, representing diastereomers produced by the reduction.^34^ Production of epoxy-enol and dihydroxy-enone analogs was also confirmed via LC-MS analysis. We detected molecular ions with *m/z* of 237 and 253, corresponding to the proton adducts of the respective analogs (Figure S1B).

**Figure 3.**
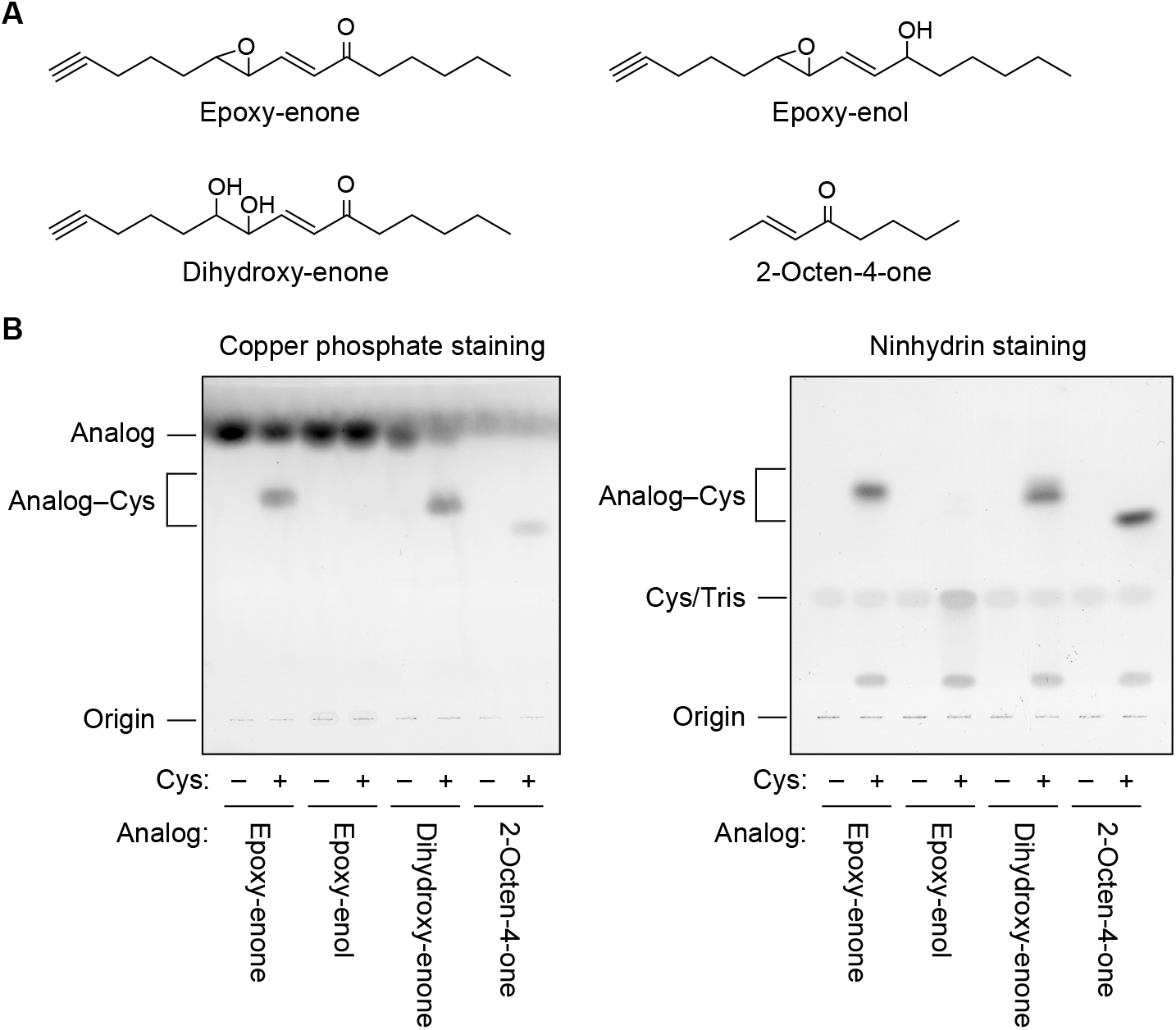
Effects of functional groups on conjugation of EE-analog with Cys. (A) Structures of EE-analog, its derivatives (epoxy-enol and dihydroxy-enone forms), and 2-octen-4-one). (B) 2 mM of each compound was mixed with 2 mM Cys, incubated at 37 °C for 1 h, separated by TLC, and visualized by copper phosphate staining (left panel) or ninhydrin staining (right panel). See also Figure S1.

Each compound was incubated with Cys, and the reaction products were separated by TLC and stained with copper phosphate and ninhydrin. Cys conjugates of EE-analog, epoxy-enol analog, and 2-octen-4-one, all of which contain enone (α,β-unsaturated carbonyl) structures, were detected by both detection methods (Figure 3B). Conversely, we observed no reaction product of the epoxy-enol analog with Cys. These results indicate that the enone structure is essential for Cys-conjugate formation, but that the epoxy group is not required.

### Formation of EE ceramide–Cys conjugate

To examine whether EE ceramides bind to Cys similarly to the EE-analog, EE ceramides were prepared from mouse epidermis. EE ceramides are known to be reversibly released from protein-bound ceramide fractions (the remnants after extraction of free lipids with organic solvents) by long-term incubation with organic solvents.^34^ The EE ceramides thus prepared were subjected to LC coupled with tandem MS (MS/MS) analysis for structural confirmation. Precursor ion scan analysis in positive ion mode for fragment ions with sphingosine structure (*m/z* = 264) detected molecular ions ([M+H]^+^ and [M+H –H_2_O]^+^) corresponding to the EE ceramides containing saturated or monounsaturated ω-OH FAs with a C30 to C36 chain length (Figures S2A and S2C). Similar results were obtained when detecting [M+HCOO]^−^ ions by precursor ion scan analysis in negative ion mode for fragment ions with the *O*-acyl chain having epoxy-enone (*m/z* = 309) (Figures S2B and S2D).

The obtained EE ceramides were incubated alone (as control) or with Cys, and the reaction products were subjected to precursor ion scan analysis to detect fragment ions with a sphingosine structure. The reaction product of EE ceramides alone produced ion peaks in the LC retention time range of 12.6 min to 13.9 min, which corresponded to the molecular ion peaks of EE ceramides with C30 to C36 ω-OH FAs (Figures 4A and 4B), as described above (Figures S2A and S2C). On the other hand, when EE ceramides and Cys were incubated, peaks were detected at 11.6 to 12.2 min preceding the LC retention time of the EE ceramide molecular ion peaks (Figure 4A, red). These peaks contained molecular ions with *m/z* of 1,190, 1,192, 1,218, and 1,246 (Figure 4B, lower panel), which corresponded to the conjugates of Cys and EE ceramides with C32:1, C32:0, C34:1, and C36:1 ω-OH FAs, respectively.

**Figure 4.**
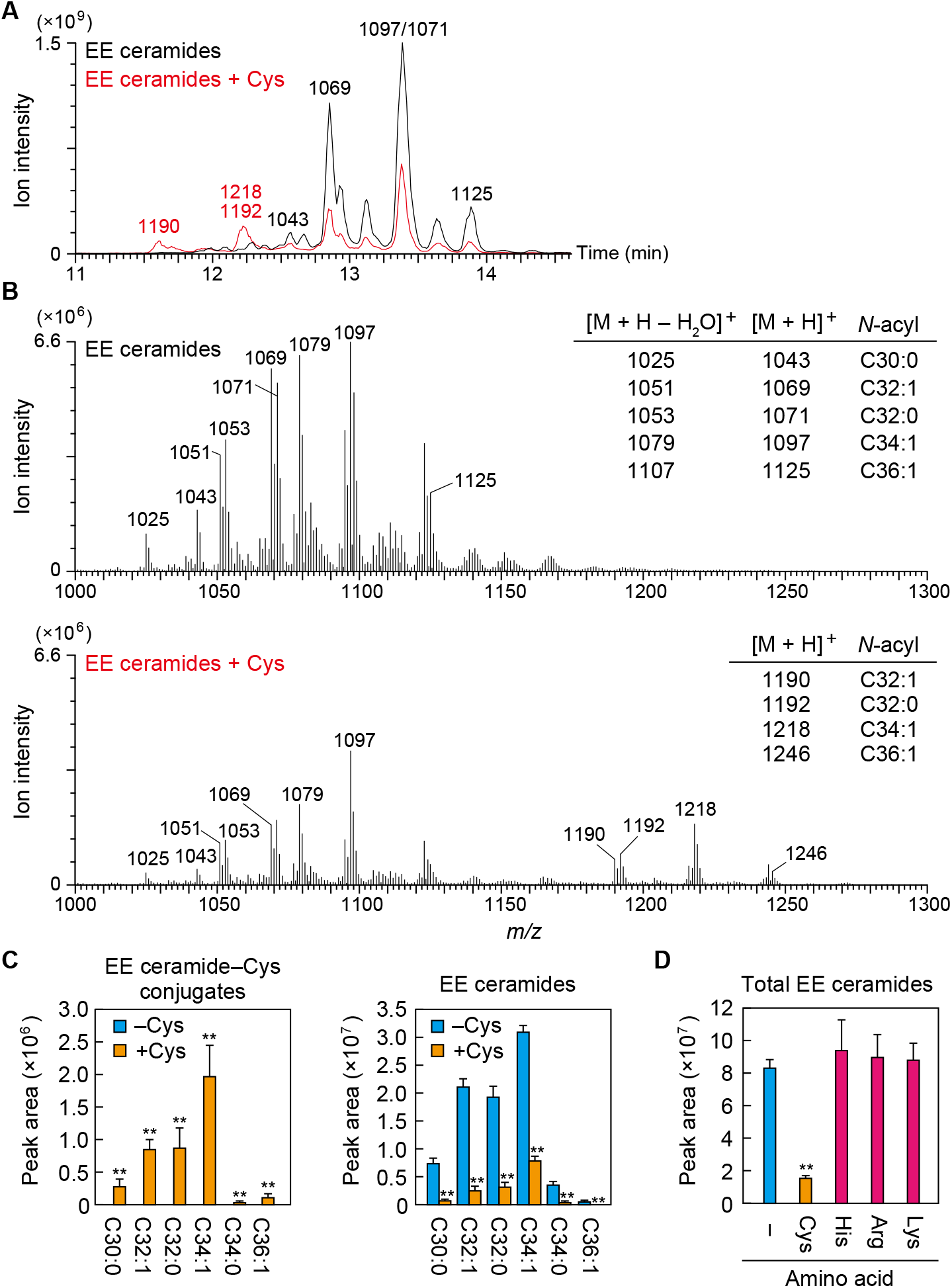
Formation of EE ceramide–Cys conjugate. (A–C) EE ceramides were mixed with 2 mM Cys and incubated at 37 °C for 1 h. (A, B) The reaction products were subjected to LC-MS/MS analysis using precursor ion scanning mode to detect precursor ions with the product ion of *m/z* = 264.3 (positive ion mode; scan range of *m/z*, 1,000–1,300). (A) Total ion current chromatogram. The *m/z* value(s) of the major precursor ions in each peak are shown above each peak. (B) Mass spectra with retention time of 11.5–14 min. The *m/z* values and *N*-acyl chains of the precursor ions corresponding to major peaks are listed on the right. (C) EE ceramide–Cys conjugates and EE ceramides were detected via LC-MS/MS analysis using MRM mode. Values represent means + SD (n = 3; Student’s *t*-test; \*\**P* < 0.01). (D) EE ceramides were mixed with an amino acid (Cys, His, Arg, or Lys; 2 mM each), incubated at 37 °C for 1 h, and subjected to LC-MS/MS analysis using MRM mode. Values represent means + SD (n = 3; Dunnett’s test vs control; \*\**P* < 0.01). See also Figures S2 and S3.

Next, we quantified EE ceramides and EE ceramide–Cys conjugates using the multiple reaction monitoring (MRM) mode of LC-MS/MS. MRM mode can detect ions specifically and with high sensitivity by selecting both precursor and product ions. Here, we selected [M+H]^+^ ions of EE ceramides or EE ceramide–Cys conjugates as the precursor ions and an ion with sphingosine structure (*m/z* = 264) as the product ion for MRM measurements. When EE ceramides and Cys were incubated, we detected EE ceramide–Cys conjugates with C30 to C36 saturated and monounsaturated ω-OH FAs (Figure 4C). The total amount of EE ceramides was reduced to 19% by the incubation with Cys compared to without Cys (Figures 4C and 4D). Conversely, when EE ceramides were incubated with His, Lys, or Arg, no molecular ions corresponding to conjugates of the respective amino acids were detected (Figure S3), with almost no change in EE ceramide levels (Figure 4D). These results indicate that EE ceramides, like the EE-analog, form stable conjugates with only Cys among the nucleophilic amino acids.

### Formation of conjugates of EE ceramides/EE-analog with Cys-containing peptides

Involucrin has been identified as the major binding target protein of P-O ceramides.^33^ Although the targets of P-EO ceramides are unknown, we examined the possibility that EE ceramides bind to involucrin using involucrin peptides. Involucrin contains two Cys residues that are conserved in mouse and human. We prepared two involucrin peptides containing these Cys residues (peptide 1, PTQCQVF; peptide 2, QKCEHQQQQ). The peptides were incubated with EE-analog and the reaction products were separated by TLC and stained with copper phosphate. We found that both peptides formed conjugates with EE-analog (Figure 5A). In contrast, the CA mutant of peptide 1 (PTQAQVF) did not form a conjugate with EE-analog. These results indicate that EE-analog binds to the peptide via Cys residue.

**Figure 5.**
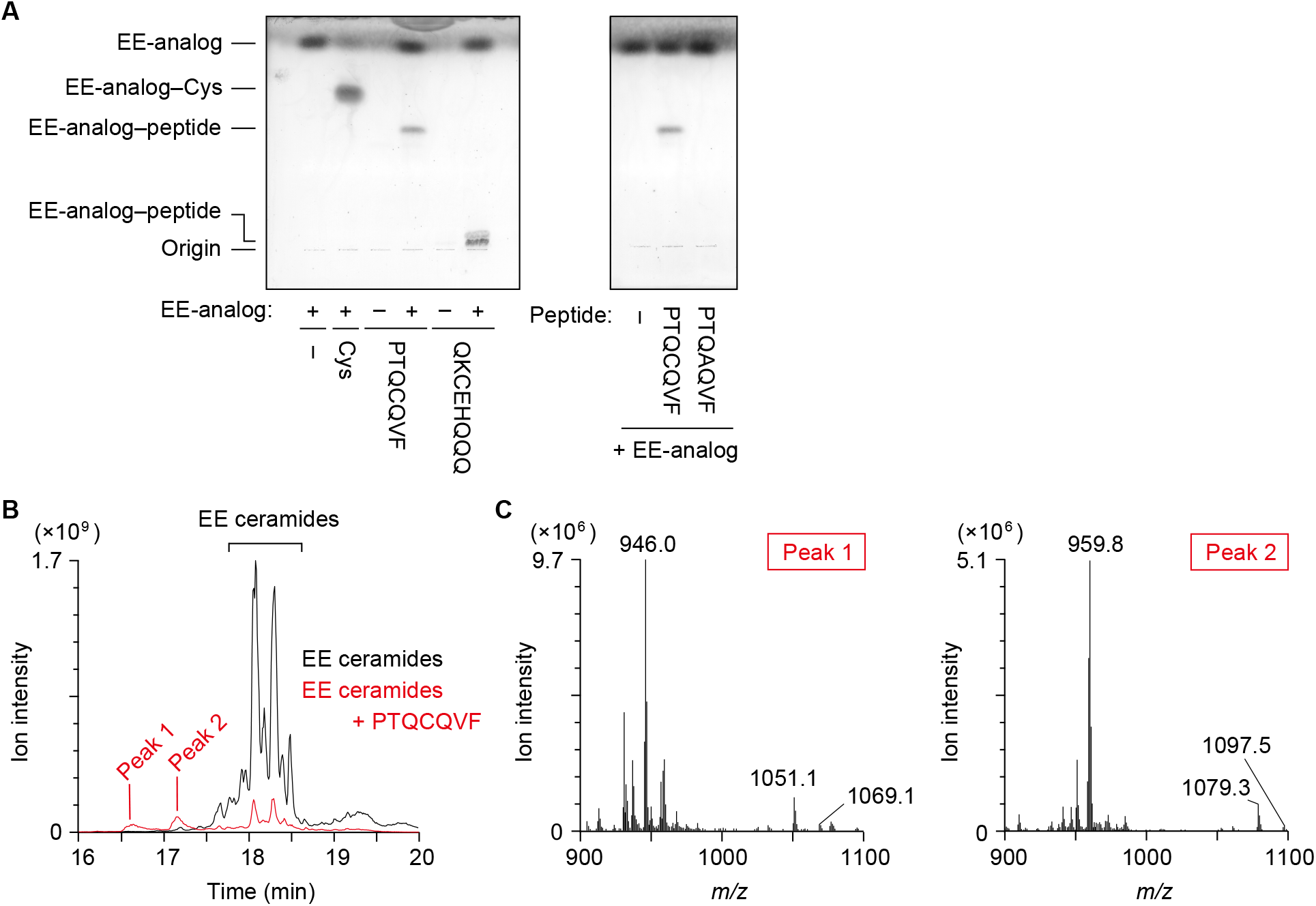
Conjugation of EE-analog/EE ceramides with Cys-containing peptides. (A) 2 mM EE-analog was mixed with 2 mM peptide (involucrin peptide 1 [PTQCQVF], peptide 2 [QKCEHQQQ], and mutant peptide 1 [PTQAQVF]), incubated at 37 °C for 1 h, separated by TLC, and visualized by copper phosphate staining. (B and C) EE ceramides were mixed with 2 mM peptide 1, incubated at 37 °C for 1 h, and subjected to LC-MS/MS analysis using precursor ion scanning mode to detect precursor ions with the product ion of *m/z* = 264.3 (positive ion mode; scan range of *m/z*, 900–1,350). Total ion current chromatogram (B) and the mass spectra of peak 1 and peak 2 (C) are shown.

Next, EE ceramides were incubated with peptide 1, and the reaction products were subjected to precursor ion scan analysis (*m/z* = 264) of LC-MS/MS. We detected two peaks (peaks 1 and 2) for the reaction products, which preceded the peaks of EE ceramides (Figure 5B). The intensities of EE ceramide peaks decreased following incubation with the peptide. Peaks 1 and 2 contained molecular ions with *m/z* = 946 and 960, respectively (Figure 5C), which corresponded to the divalent ions of the conjugates of peptide 1 ([M+H]^+^; *m/z* = 822) and EE ceramides containing C32:1 and C34:1 ω-OH FAs, respectively ([M+H]^+^; peak 1, *m/z* = 1,069; peak 2, *m/z* = 1,097). These results indicate that EE ceramides can bind at least to involucrin peptides.

### Presence of Cys-bound P-EO ceramides in epidermis

To examine whether Cys-bound P-EO ceramides are indeed present in epidermis, protein-bound ceramide fraction of mouse epidermis was treated with pronase E to degrade proteins to amino acids, followed by precursor ion analysis (*m/z* = 264) by LC-MS/MS. We detected Cys-bound P-EO ceramides with C30:0 (*m/z* = 1,164), C32:1 (*m/z* = 1,190), C32:0 (*m/z* = 1,192), and C34:1 (*m/z* = 1,218) ω-OH FAs in wild type (WT) epidermis in a pronase E treatment-dependent manner (Figure 6A). These peaks were not detected in the epidermis of KO mice for the FA ω-hydroxylase *Cyp4f39* (Figure 6B), which cannot produce protein-bound ceramides.^13^ MRM analysis of LC-MS/MS also confirmed the presence of Cys-bound P-EO ceramides in WT mice and the absence in *Cyp4f39* KO mice (Figure 6C). No molecular ions corresponding to the conjugates of EE ceramides to other nucleophilic amino acids, such as His, Arg, or Lys, were detected in the epidermis of WT mice (Figure 6A). The predicted *m/z* values of Michael addition reaction products of EE ceramides with C34:1 ω-OH FA are 1,252 (His), 1,271 (Arg), and 1,243 (Lys), and those of Schiff base products are 1,253 (Arg) and 1,225 (Lys).

**Figure 6.**
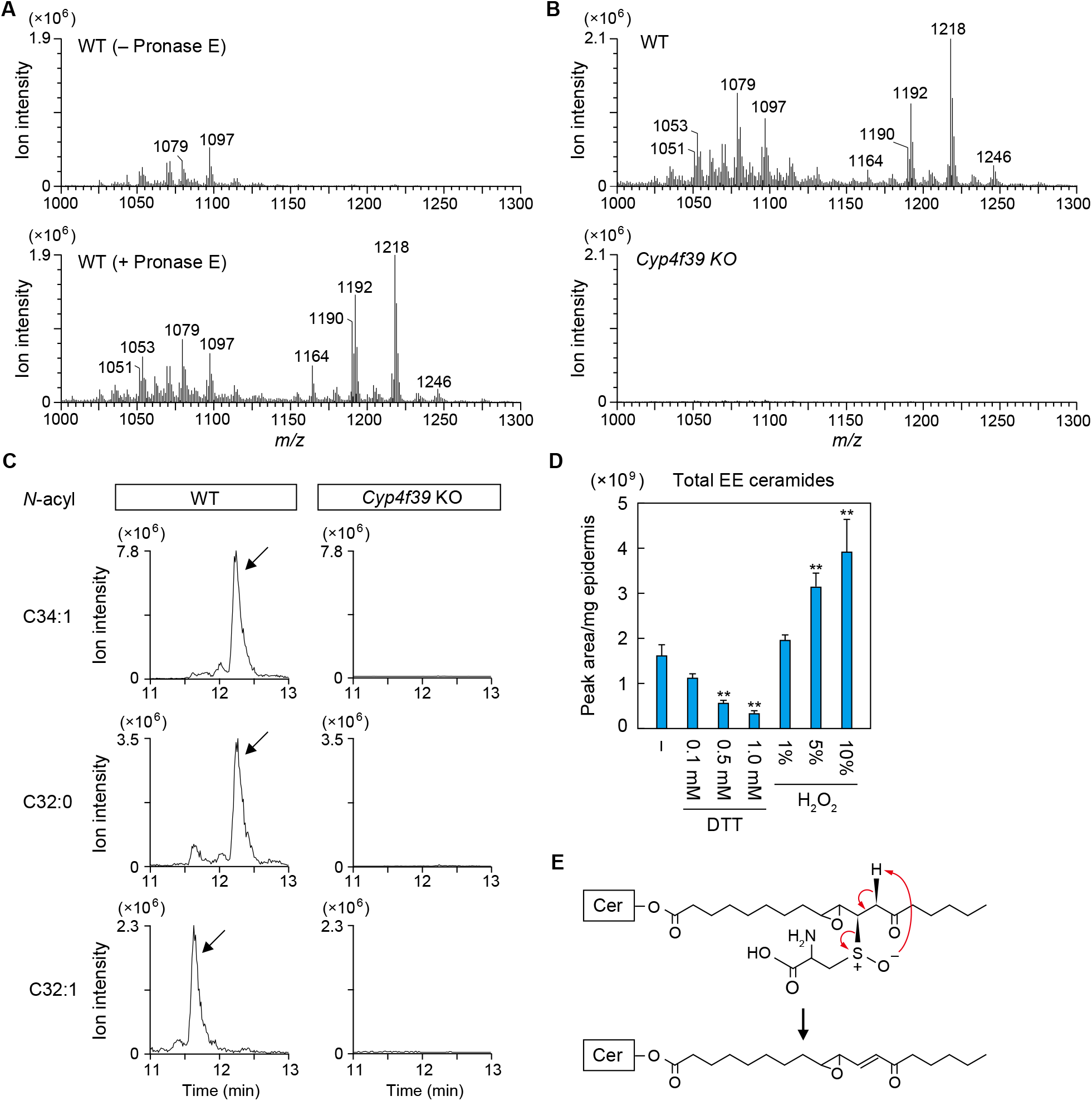
Presence of Cys-bound P-EO ceramides in the epidermis. (A–C) Protein-bound ceramide fractions prepared from WT and *Cyp4f39* KO mouse epidermis (10 mg) were digested with pronase E (1 mg/mL) at 37 °C for 2 h. (A and B) Digested samples were subjected to LC-MS/MS analysis using precursor ion scanning mode to detect precursor ions with the product ion of *m/z* = 264.3 (positive ion mode; scan range of *m/z*, 1,000–1,300). The mass spectra at the retention time during which Cys–P-EO ceramides are eluted are shown. (C) Cys-bound P-EO ceramides containing C34:1, C32:0, or C32:1 as an *N*-acyl moiety were detected via LC-MS/MS analysis using MRM mode. Cys-bound P-EO ceramide peaks are indicated by arrows. (D) Protein-bound ceramide fraction prepared from WT mouse epidermis (0.05 mg) was incubated at 60 °C for 3 h in 90% CH_3_OH containing DTT or H_2_O_2_ with the indicated concentrations. EE ceramides released from protein-bound ceramide fraction were detected via LC-MS/MS analysis using MRM mode. Values represent means + SD (n = 3; Dunnett’s test vs. no addition; \*\**P* < 0.01). (E) Mechanism of sulfoxide elimination. Oxidation of the sulfur atom that constitutes the thioether linkage between EE ceramides and Cys residue accelerates the release of EE ceramides. See also Figure S4.

We next examined the presence of P-O ceramides in the protein-bound ceramide fraction of the pronase E-treated WT epidermis using MRM analysis. Although retention times of P-O ceramides were predicted to be shorter than those of ω-OH ceramides (∼12.0 min, ω-OH ceramide C34:1), no peaks were detected in these regions (Figure S4). We detected pronase E-dependent peaks at around 13 min with the MRM setting for P-O ceramide with C34:1 ω-OH FA. However, as the retention times were slower than expected, these peaks may not represent P-O ceramide. In summary, we detected only Cys-bound P-EO ceramides as protein-bound ceramides in the epidermis of WT mice.

It is known that the cleavage of the thioether bond between the β-carbon of the carbonyl group and the thiol, i.e., the release of α,β-unsaturated carbonyl compounds, is stimulated by oxidation of sulfur atoms (sulfoxide elimination).^39^ We next investigated whether the reversibly eluted EE ceramides from the protein-bound ceramide fraction are generated via sulfoxide elimination of Cys-bound P-EO ceramides. For this purpose, a reducing agent (dithiothreitol; DTT) or an oxidizing agent (H_2_O_2_) was added during the incubation of the protein-bound ceramide fraction with organic solvent (90% CH_3_OH). Elution of EE ceramides from the protein-bound ceramide fraction decreased in a DTT concentration-dependent manner (0.1 mM, 70%; 0.5 mM, 35%; 1 mM, 21%), while addition of H_2_O_2_ increased elution (1%, 1.2 fold; 5%, 2.0 fold; 10%, 2.4 fold) (Figure 6D). These results suggest that EE ceramides are released from Cys-bound P-EO ceramides via sulfoxide elimination (Figure 6E).

### Abundance of Cys-bound P-EO ceramides in the epidermis

We performed quantitative analysis to determine the proportion of P-EO ceramides in protein-bound ceramides in epidermis. The protein-bound ceramide fraction (total protein-bound ceramides), reversibly bound fraction (Cys-bound P-EO ceramides), and irreversibly bound fraction (protein-bound ceramides containing P-O ceramides, with possible others) were each converted to ω-OH ceramides by alkaline treatment, and the resulting ω-OH ceramides were quantified by LC-MS/MS (Figure 7A). When preparing the reversibly bound fraction, we repeated the elution procedure five times, and each elution was measured. Fourth and fifth elution fraction levels were lower than the first three elution fractions (Figure 7B), suggesting that most reversible protein-bound ceramides were eluted by five elutions. The amount of reversible protein-bound ceramides (total amount from 1–5 elution fractions) and irreversible protein-bound ceramides was 60% and 46% of the total amount of protein-bound ceramides, respectively (Figure 7C). Although it is unknown how much P-EO ceramide remains un-eluted in the irreversibly bound fraction, these results indicate that Cys-bound P-EO ceramides are the predominant protein-bound ceramides.

**Figure 7.**
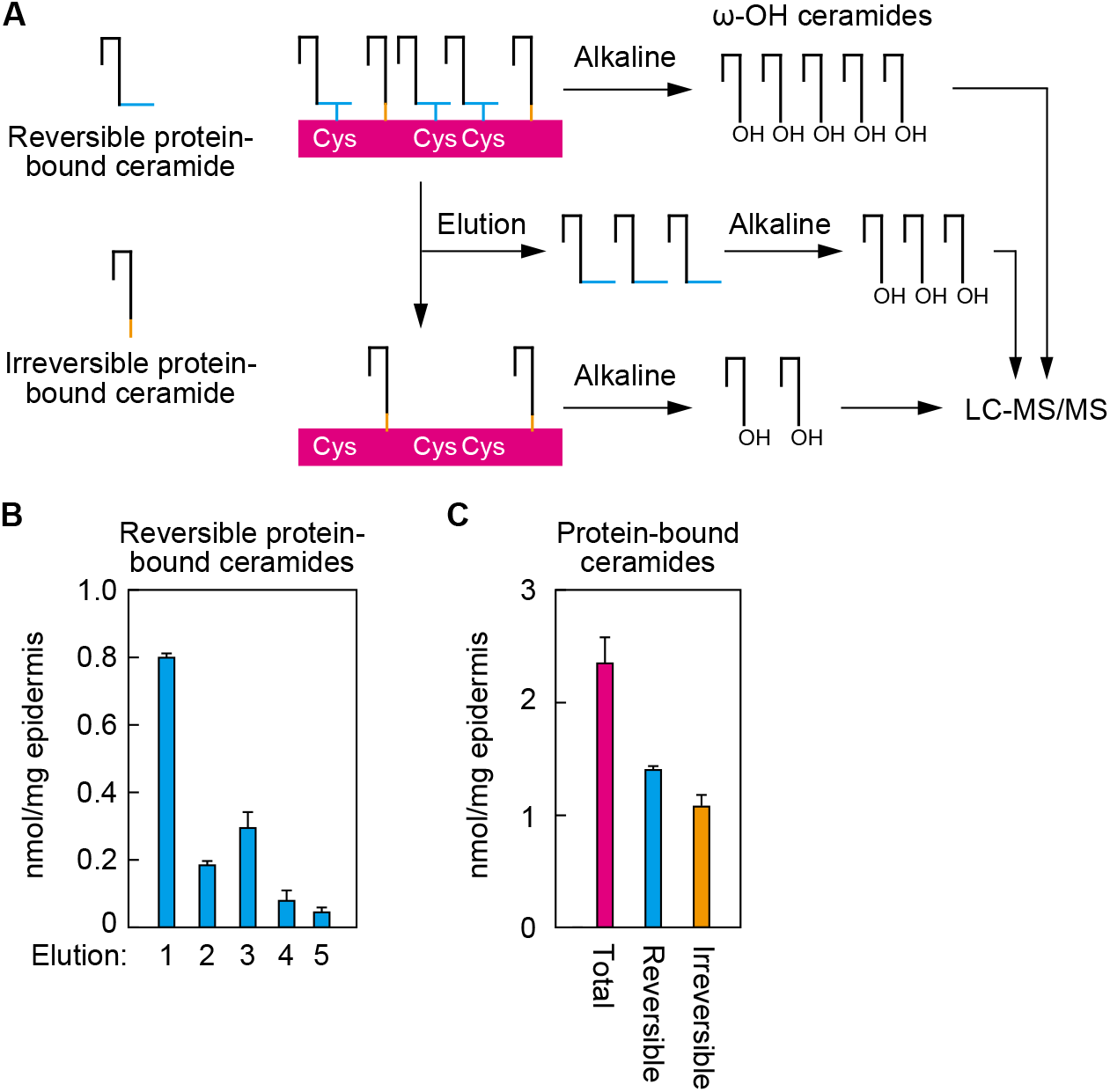
Abundance of reversible protein-bound ceramides in the epidermis. (A) Schematic illustration of preparation and detection of total, reversible, and irreversible protein-bound ceramides. (B and C) The protein-bound ceramide fraction was incubated with CHCl_3_/CH_3_OH = 1:2 (v/v) at 60 °C five times (first two times for 3 h; third time for 16 h; last two times for 3 h). The protein-bound ceramide fraction (total protein-bound ceramides; C), each elution fraction (B), the mixture of five elution fractions (reversible protein-bound ceramides; C), and the pellet fraction after five elutions (irreversible protein-bound ceramides; C) were treated with alkaline to release ω-OH ceramides, followed by LC-MS/MS analysis. Values represent means + SD (n = 3).

## Discussion

Although the P-O ceramide model has long been considered the structure model of protein-bound ceramides,^33^ the P-EO ceramide model has been recently introduced as an alternative.^31^ The P-EO model predicts that the enone moiety of EE ceramides binds to nucleophilic amino acid residues of the CE proteins by Schiff base formation or the Michael addition reaction. In this study, we found that P-EO ceramides are specifically formed by the Michael addition reaction of EE ceramides to Cys residues (Figures 4, 6, S3). According to the HSAB rule (hard and soft acids and bases principle), which classifies the reactivity of functional groups, the β-carbon of α,β-unsaturated carbonyl belongs to the soft electrophile group and is highly reactive with soft nucleophiles, including thiol groups.^40,41^ The amino group is classified as a hard nucleophile and the imidazole group of His is an intermediate nucleophile. Therefore, their reactivity to β-carbon is low, which means that forming a bond with EE ceramides is difficult. Carbonyl carbon is a hard electrophile and is generally highly reactive with amino groups, forming an imine. Since this reaction is reversible in the presence of water, it is possible that imine was cleaved during our extraction procedure.

In the experiment that led to the development of the P-O ceramide model, peptide–protein-bound ceramide conjugates obtained from the protein-bound ceramide fraction of human foreskin by proteinase K treatment were analyzed via MS.^33^ However, only six peaks of peptide–protein-bound ceramide conjugates were analyzed, which is estimated to have covered 15–20% of the total protein-bound ceramides. In the present study, we could not detect any peaks corresponding to P-O ceramide (Figure S3). These results suggest the possibility that P-O ceramides are minor components of protein-bound ceramides or even artifacts under certain experimental conditions.

In the predicted P-O ceramide synthesis pathway, a bond is formed between the ω-OH ceramides and CE proteins in the final step. Under *in vitro* conditions at least, the transglutaminase TGM1 exhibited this bond-forming activity.^35^ *In vivo*, protein-bound ceramide levels in the epidermis of *Tgm1* KO mice were ∼60% of those in WT mice.^42^ Since TGM1 is involved in CE formation by catalyzing protein cross-linking, this reduction may be an indirect effect of impaired CE formation. In the present study, we showed that approximately 60% of protein-bound ceramides bind reversibly to CE proteins (Figure 7C). This suggests that at least 60% of the protein-bound ceramides are Cys-bound P-EO ceramides. The exact molecular nature of the protein-bound ceramides in the remaining ∼40% that were present in the irreversibly bound fraction is unknown. These could be either P-O ceramides or irreversible forms of P-EO ceramides, in which the epoxy or keto group may further react with other amino acids of CE proteins.

In the binding assays with Cys using EE-analog and their derivatives, the epoxy group did not affect binding of the thiol to the β-carbon of enone (Figure 3). The epoxy group also did not affect enone formation (reduction of the carbonyl carbon) by SDR9C7 *in vitro.*^31^ The exact role of the epoxy group in EE ceramides/P-EO ceramides is therefore unknown, but the following two possibilities may be considered. First, the epoxy group is involved in forming a bond with other amino acids after the production of Cys-bound P-EO ceramides, as described above. This further binding would cause P-EO ceramides to lose their reversibility and stabilize P-EO ceramides. Second, it is possible that the epoxy group of EE ceramide promotes the reaction of the β-carbon of the enone moiety with Cys residues *in vivo*. For example, the epoxy group can bring EE ceramides into close proximity with Cys residues by interacting with polar groups of CE proteins. Alternatively, the epoxy group may itself promote reactivity between EE ceramides and the thiol group of Cys residues. The p*K*_a_ of the thiol group of the Cys residue is 8.4 and the pH of the stratum corneum is 4–6.^43^ The thiol group of the CE proteins is thus protonated and consequently not highly nucleophilic. However, it is possible that the unshared electron pair of the oxygen atom of the epoxy group attracts the thiol group proton, thereby enhancing thiol nucleophilicity and promoting the Michael addition reaction between the thiol group and the EE ceramides.

In this study, we selected the peptides of involucrin, a major target of P-O ceramides,^33,35^ for the binding assay with EE-analog and EE ceramides (Figure 5). Two Cys residues in involucrin are conserved between humans and mice, and both of the Cys-containing peptides were found to bind to EE-analog/EE ceramides in the binding assay. Desmoplakin, envoplakin, and periplakin in addition to involucrin have been identified as CE proteins that bind to P-O ceramides.^33^ Since these proteins have more conserved Cys residues in humans and mice (40, 21, and 8 Cys residues, respectively) than involucrin, it is possible that these are more predominant targets of P-EO ceramides than involucrin.

In this study, we revealed that the structure of protein-bound ceramides is that of Cys-bound P-EO ceramides. Furthermore, we established a method to detect these separately from P-O ceramides. Our findings and analytical method can help develop future diagnosis of skin diseases caused by skin barrier abnormalities, and therapeutic agents relevant to such diseases.

## Limitations of the study

In the present study, we revealed that the target amino acid residue of P-EO ceramides is Cys, but did not yet identify the target proteins. In addition, the exact structure of the protein-bound ceramides in the irreversibly bound fraction, which may be that of P-O ceramides or P-EO ceramides other than the Cys-bound form, is still unknown. These unsolved problems need to be clarified in future research.

## Acknowledgements

This work was supported by funding from the Takeda Science Foundation (to A.Kihara), by Japan Society for the Promotion of Science (JSPS) KAKENHI grant numbers JP22K08372 (to Y.O.), JP22K15241 (to A.Katsuyama), JP22H02738 (to I.S.), and JP22H04986 (to A.Kihara), and by a grant from the Akiyama Life Science Foundation (to Y.O.). NMR analyses were performed using JNM-ECZ400, which is registered in the open facility managed by the Global Facility Center, Creative Research Institution, Hokkaido University.

## Author contributions

Conceptualization, Y.O. and A.Kihara; Methodology; Y.O.; Investigation, Y.O., T.N., T.I., and A.Katsuyama; Writing – Original Draft, Y.O., A.Katsuyama, and A.Kihara; Writing – Review & Editing, A.Kihara; Supervision, S.I. and A.Kihara; Project Administration, A.Kihara; Funding Acquisition, Y.O. A.Katsuyama., S.I., and A.Kihara.

## STAR METHODS

## KEY RESOURCES TABLE

Refer to a separate Word file.

## RESOURCE AVAILABILITY

### Lead contact

Further information and requests for resources and reagents should be directed to and will be fulfilled by the Lead Contact, Akio Kihara.

### Materials availability

All unique/stable reagents generated in this study are available from the Lead Contact with a completed Materials Transfer Agreement.

### Data and code availability

- All data generated in this study are included in the manuscript.
- This paper does not report original code.
- Any additional information required to reanalyze the data reported in this paper is available from the lead contact upon request.

## EXPERIMENTAL MODEL AND SUBJECT DETAILS

### Mice

C57BL/6J mice were purchased from Sankyo Labo Service Corporation (Tokyo, Japan). *Cyp4f39* KO mice were previously generated using C57BL/6J background.^13^ Mice were maintained under specific pathogen-free conditions at a room temperature of 23 °C ± 1 °C, at 50 ± 5% humidity under a 12-hour light-dark cycle, and provided *ad libitum* access to water and food (CLEA Rodent Diet CE-2; CLEA Japan, Tokyo, Japan). Animal experiments were approved by the institutional animal care and use committee of Hokkaido University and conducted in accordance with their guidelines.

### Cell culture and transfection

HEK 293T cells were obtained from RIKEN BioResource Research Center (Tsukuba, Japan) and cultured in Dulbecco’s Modified Eagle’s medium (D6429; Merck, Darmstadt, Germany) supplemented with 10% fetal bovine serum (Thermo Fisher Scientific), 100 units/mL penicillin, and 100 μg/mL streptomycin (Merck). Cells were grown on collagen coat dishes (IWAKI, Shizuoka, Japan) and incubated under 5% CO_2_ at 37 °C. Transfection was performed using PEI Max (Polysciences, Warrington, UK) as described previously.^44^

## METHOD DETAILS

### Synthesis of EE-analog

EE-analog was synthesized via a scheme composed of six reaction steps (Figure S5), where each compound was confirmed by NMR spectrometry and MS analysis. Compound 1 (ethyl (*E*)-oct-2-en-7-ynoate) was synthesized as below. 5-Hexyn-1-ol (500 μL, 4.53 mmol; Tokyo Chemical Industry, Tokyo, Japan) was added dropwise to a mixture of Dess-Martin periodinane (2.50 g, 5.89 mmol; Tokyo Chemical Industry) in CH_2_Cl_2_ (45 mL) at 0 °C over 20 min. The mixture was warmed to room temperature and stirred for 2.5 h. The reaction was quenched with a solution of *sat*. *aq*. NaHCO_3_ and *sat*. *aq*. Na_2_S_2_O_3_, after which the mixture was partitioned between CH_2_Cl_2_ and *sat*. *aq*. NaHCO_3_. The organic phase was washed with *sat*. *aq*. NaHCO_3_ and brine, then Na_2_SO_4_ was added to the organic phase to remove trace amounts of water. The mixture was filtered and concentrated *in vacuo* to afford a crude aldehyde. A mixture of the crude aldehyde and triethyl phosphonoacetate (988 μL, 4.98 mmol; Tokyo Chemical Industry) was treated with 1,8-diazabicyclo[5.4.0]undec-7-ene (DBU; 1.02 mL, 6.80 mmol; Tokyo Chemical Industry) at 0 °C for 16 h. The mixture was partitioned between ethyl acetate and 1 M *aq*. HCl. The organic phase was washed with 1 M *aq*. HCl, *sat*. *aq*. NaHCO_3_ (× 2), and brine, dried (Na_2_SO_4_), filtered, and concentrated *in vacuo*. The residue was then purified by flash silica gel (Silica gel 60N; Kanto chemical, Tokyo, Japan) column chromatography (ϕ 2 cm × 20 cm, 0–5% ethyl acetate/hexane) to afford compound 1 (320 mg, 2.10 mmol, 46%) as a colorless oil. ^1^H NMR (CDCl_3_; Cambridge Isotope Laboratories, Tewksbury, MA, USA; 400 MHz) δ 6.94 (dt, 1H, *J* = 15.6, *J* = 7.0 Hz), 5.86 (dt, 1H, *J* = 15.6, *J* = 1.4 Hz), 4.18 (q, 2H, *J* =7.0 Hz), 2.33 (tdt, 2H, *J* = 7.3 Hz, *J* = 7.0, *J* = 1.4 Hz), 2.23 (td, 2H, *J* =7.3, *J* = 2.7 Hz), 1.97 (t, 1H, *J* = 2.7 Hz), 1.69 (tt, 2H, *J* = 7.3 Hz), 1.29 (t, 3H, *J* =7.0 Hz). This is a known and reported compound.^45^

Compound 2 ((*E*)-oct-2-en-7-yn-1-ol) was synthesized as below. A solution of compound 1 (320 mg, 1.93 mmol) in CH_2_Cl_2_ (15 mL) was treated with diisobutylaluminium hydride (1 M in toluene, 4.24 mL, 4.24 mmol; Tokyo Chemical Industry) at –78 °C for 3 h. The mixture was warmed to room temperature and stirred for 11 h. The reaction was quenched with *sat. aq.* NH_4_Cl at 0 °C. After stirring the mixture at room temperature for several minutes, an insoluble material was filtered off through a Celite pad (Merck) and the filtrate partitioned between CH_2_Cl_2_ and brine. The organic phase was washed with brine, dried (Na_2_SO_4_), filtered, and concentrated *in vacuo*. The residue was purified by flash silica gel column chromatography (ϕ 2 cm × 20 cm, 34% ethyl acetate/hexane) to afford compound 2 (149.7 mg, 1.21 mmol, 62%) as a colorless oil. ^1^H NMR (CDCl_3_, 400 MHz) δ 5.68–5.67 (m, 2H), 4.10 (br s, 2H), 2.31–2.15 (m, 4H), 1.96 (t, 1H, *J* = 2.8 Hz), 1.62 (tt, 2H, *J* = 7.2 Hz); *m/z* 124.9 [M+H]^+^.

Compound 3, ((2*R*,3*R*)-3-(pent-4-yn-1-yl)oxiran-2-yl)methanol, was synthesized as below. A solution of compound 2 (50 mg, 0.40 mmol) in CH_2_Cl_2_ was added to a suspension of diisopropyl D-tartrate (10.9 μL, 0.052 mmol; Tokyo Chemical Industry), tetraisopropyl orthotitanate (10 mg, 0.035 mmol; Tokyo Chemical Industry), and molecular sieves 4A (20 mg; Nacalai Tesque, Kyoto, Japan) at –20 °C. We then added *tert*-butyl hydroperoxide (5.5 M in decane; Merck) dropwise for 1 min, and the mixture was stirred at –20 °C for 2 h. The reaction was quenched with triphenylphosphine (Tokyo Chemical Industry), then purified by flash silica gel column chromatography (<φ 1 cm × 10 cm, 0–20–40% ethyl acetate/hexane) to produce compound 3 (33.1 mg, 0.24 mmol, 59%) as a colorless oil. ^1^H NMR (CDCl_3_, 400 MHz) δ 3.91 (dd, 1H, *J* = 12.4, *J* = 2.5 Hz), 3.64 (dd, 1H, *J* = 12.4, *J* = 4.1 Hz), 2.98 (ddd, 1H, 1H, *J* = 2.5, *J* = 4.6 Hz), 2.95 (ddd, 1H, 1H, *J* = 2.5, *J* = 4.1 Hz), 2.28–2.25 (m, 2H), 1.97 (t, 1H, *J* = 2.7 Hz), 1.80–1.64 (m, 5H); *m/z* 163.2 [M+Na]^+^.

EE-analog ((*E*)-1-((2*R*,3*R*)-3-(pent-4-yn-1-yl)oxiran-2-yl)oct-1-en-3-one) was synthesized as below. A mixture of compound 3 (15 mg, 0.12 mmol) and NaHCO_3_ (12.9 mg, 0.15 mmol) in CH_2_Cl_2_ (5 mL) was treated with Dess-Martin periodinane (65 mg, 0.15 mmol) at 0 °C for 5 min. The mixture was warmed to room temperature and stirred for 16.5 h. The reaction was quenched with a solution of *sat*. *aq*. NaHCO_3_ and *sat*. *aq*. Na_2_S_2_O_3_, then the mixture was partitioned between ethyl acetate and *sat*. *aq*. NaHCO_3_. The organic phase was washed with brine, dried (Na_2_SO_4_), filtered, and concentrated *in vacuo* to produce a crude aldehyde. A mixture of the crude aldehyde in CH_2_Cl_2_ was treated with compound 4 (57.3 mg, 0.15 mmol), which was synthesized as reported ^46^ at 0 °C for 19 h. The mixture was purified by scraping from a TLC plate (Silica gel 60 F254 TLC plates, Merck) and eluting (25% ethyl acetate/hexane) to afford EE-analog (6.0 mg, 0.026 mmol, 21% over 2 steps) as a colorless oil. ^1^H NMR (CDCl_3_, 400 MHz) δ 6.51 (dd, 1H, *J* = 15.9, *J* = 6.9 Hz), 6.39 (d, 1H, *J* = 15.9 Hz), 3.24 (dd, 1H, *J* = 6.4, *J* = 1.8 Hz), 2.92 (td, 1H, *J* = 5.0, *J* = 1.8 Hz), 2.52 (t, 2H, *J* = 7.3 Hz), 2.27 (td, 2H, *J* = 6.4, *J* = 2.7 Hz), 1.97 (t, 1H, *J* = 2.7 Hz), 1.86–1.66 (m, 4H), 1.61 (dd, 2H, *J* = 7.3 Hz), 1.37–1.25 (m, 4H), 0.89 (t, 3H, *J* = 7.1 Hz); *m/z* 235.1 [M+H]^+^.

### Preparation of the epoxide hydrolase

The plasmid to express epoxide hydrolase EPHX2 in mammalian cells was generated as follows. The coding sequence of *EPHX2* was amplified by PCR from liver cDNA (Human Multiple Tissue cDNA panels; Takara Bio, Shiga, Japan) using primers (5’-<u>GTCGAC</u>ATGACGCTGCGCGCGGCCGTCTTCG-3’, 5’-CTACATCTTTGAGACCACCGGTGGG-3’; *Sal*I sites underlined) and cloned into the TA cloning vector pGEM-T Easy Vector (Promega, Madison, WI, USA). The *EPHX2* fragment was excised with restriction enzymes *Sal*I and *Not*I and cloned into the pCE-puro 3×FLAG-1, the mammalian expression vector to produce N-terminally 3×FLAG-tagged proteins,^47^ yielding pCE-puro 3×FLAG-EPHX2.

EPHX2-containing soluble fraction was prepared from HEK 293T cells overexpressing 3×FLAG-EPHX2 as follows. After 24 h from transfection of pCE-puro 3×FLAG-EPHX2 into HEK 293T cells, the cells were washed with PBS, harvested by a scraper, and transferred to a plastic tube. The cells were suspended in lysis buffer [50 mM sodium phosphate buffer (pH 7.4), 0.1 mM EDTA, 1 mM phenylmethylsulfonyl fluoride (FUJIFILM Wako Pure Chemical Corporation; Osaka, Japan), 1× protease inhibitor cocktail (cOmplete, EDTA-free protease inhibitor cocktail; Merck)], and lysed by sonication. Unbroken cells were removed by centrifugation (400 × *g*, 4 °C, 3 min) and the total cell lysates thus prepared were subjected to ultracentrifugation (100,000 × *g*, 4 °C, 35 min). The resulting supernatant (soluble fraction) was recovered and used as an enzyme source of dihydroxy-enone-analog preparation, as described below.

### Synthesis of EE-analog derivatives

Epoxy-enol-analog was synthesized by reducing the EE-analog. 10 mM EE-analog in 150 µL of CH_3_OH were incubated with 2 mg/mL NaBH_4_ (Merck) in 150 µL of CH_3_OH at 37 °C for 1 h. Following the reactions, samples were mixed with 300 µL of CHCl_3_ and 270 µL of H_2_O and centrifugated (20,400 × *g*, room temperature, 3 min). The lower, organic phase containing epoxy-enol-analog was recovered, dried, and dissolved in 750 µL of CH_3_OH.

Dihydroxy-enone-analog was produced by cleaving the epoxy group using the soluble fraction prepared from HEK 293T cells overexpressing EPHX2, as described above. The mixture of 20 µL of 10 mM EE-analog and 180 µL of the soluble fraction (100 µg protein) was incubated at 37 °C for 1 h. Samples were then mixed with 750 µL of CHCl_3_/CH_3_OH=1:2 (v/v), 250 µL of CHCl_3_, and 250 µL of H_2_O, successively. After centrifugation (20,400 × *g*, room temperature, 3 min), the lower, organic phase containing dihydroxy-enone analog was recovered, dried, and dissolved in 100 µL of CH_3_OH.

The structures of the produced compounds were confirmed by TLC and LC-MS analysis. EE-analog and its derivatives were spotted onto a TLC plate (silica gel 60 high-performance TLC plates; Merck) and separated using CHCl_3_/CH_3_OH/acetic acid = 94:4:1 (v/v) as the solvent system. The plate was sprayed with copper phosphate solution [8% (v/v) aqueous phosphoric acid containing 3% (w/v) copper acetate], dried, and heated at 180 °C for 5 min. The LC-MS analysis is described below.

### Binding assay using EE-analog

The mixture of 5 µL of 2 mM EE-analog in CH_3_OH and 5 µL of 2 mM amino acid (Cys, Ser, His, Arg, or Lys) in 50 mM Tris-HCl (pH 7.4) were incubated at 37 °C for 1 h. The reaction products were spotted onto TLC plate and separated using 1-butanol/acetic acid/H_2_O = 3:1:1 (v/v) as the solvent system. The TLC plate was sprayed with copper phosphate solution or ninhydrin solution (0.3% [w/v] ninhydrin dissolved in 1-butanol/acetic acid [100:3, v/v]), dried, and heated at 180 °C for 5 min (copper phosphate solution) or at 120 °C for 5 min (ninhydrin staining). The binding assays using EE-analog derivatives and Cys, or EE-analog and peptides, were also conducted using the same procedure.

The produced EE-analog–Cys conjugate was extracted for LC/MS analysis as follows. A portion of the above reaction product (10 µL) was mixed with 295 µL of CH_3_OH, 300 µL of CHCl_3_, and 265 µL of H_2_O, successively. After centrifugation (20,400 × *g*, room temperature, 3 min), the lower, organic phase was collected, dried, dissolved in 1 mL CH_3_OH, and subjected to LC-MS analysis (injection volume, 5 µL), as described below.

### _1_H NMR spectrometry of EE-analog and EE-analog–Cys

The mixture of 250 µL of 44 mM EE-analog (dissolved in CD_3_OD; Cambridge Isotope Laboratories) and 250 µL of 112 mM Cys (dissolved in D_2_O [Cambridge Isotope Laboratories] containing 112 mM triethylamine) was incubated at 25 °C for 5 min and subjected to ^1^H NMR spectrometry at 400 MHz (JNM-ECZ400; JEOL, Tokyo, Japan).

### Preparation of EE ceramides

Skin was collected from mice sacrificed by decapitation at postnatal day 0 and incubated with 2 mg/mL Dispase II (Thermo Fisher Scientific, Waltham, MA, USA) in PBS at 4 °C for 16 h. The epidermis was separated from the dermis using forceps, washed with PBS, and weighed after removing excess water by pressing onto filter paper. The epidermis (5 mg) was transferred into a tube containing zirconia beads (ϕ1 mm; TOMY Seiko, Tokyo, Japan), suspended in 500 μL CHCl_3_/CH_3_OH (1:2, v/v), and crushed (4,500 rpm, 4 °C, 1 min) using a Micro Smash MS-100 (TOMY Seiko). The solutions were transferred to a glass tube and centrifugated (2,600 × *g*, room temperature, 3 min), and supernatants were removed. The pellets were then treated with 2 mL of CHCl_3_/CH_3_OH (1:2, v/v), vigorously mixed, and centrifuged (2,600 × *g*, room temperature, 3 min). To remove free lipids completely, this procedure was repeated five times, and the resulting pellets were recovered as protein-bound ceramide fractions.

EE ceramides were prepared by repeating the following procedure three times. The protein-bound ceramide fraction was incubated with 500 µL of CHCl_3_/CH_3_OH (1:2, v/v) at 60 °C for 3 h (first and second rounds) or for 16 h (third round) and centrifuged (20,400 × *g*, room temperature, 3 min) to separate supernatants and pellets. The supernatants were collected, and 300 µL of CHCl_3_/CH_3_OH (1:2, v/v) was added to the pellets each time. The supernatants were then pooled, dried, dissolved in 500 µL of 90% CH_3_OH, and mixed with 500 µL of hexane. After centrifugation (20,400 × *g*, room temperature, 3 min), the upper phase was collected. To the lower phase, 500 µL of hexane was added again and vigorously mixed. After centrifugation (20,400 × *g*, room temperature, 3 min), the upper phase was collected. The two upper phases were combined, dried, and dissolved in CH_3_OH.

In the experiment to determine whether the reversible release of EE ceramides was due to sulfoxide elimination, the protein-bound ceramide fractions (0.05 mg) were incubated at 60 °C for 3 h with 300 µL of 90% CH_3_OH containing DTT or H_2_O_2_. After centrifugation (20,400 × *g*, room temperature, 3 min), the supernatants were collected, mixed with 270 µL of CHCl_3_ and 213 µL of H_2_O, successively, and centrifuged again (20,400 × *g*, room temperature, 3 min). The lower, organic phase was collected, dried, dissolved in 1.5 mL of CHCl_3_/CH_3_OH (1:2, v/v), and subjected to LC-MS/MS analysis (injection volume, 5 µL) as descried below.

### Separation and quantification of reversible and irreversible protein-bound ceramides

The protein-bound ceramide fractions prepared from 5 mg of the epidermis as described above were suspended in 1 mL CH_3_OH, after which 10 µL of the suspension (corresponding to 0.05 mg of the epidermis) was aliquoted. The following procedure was repeated five times to elute reversible protein-bound ceramides. Samples were incubated with 300 µL of CHCl_3_/CH_3_OH (1:2, v/v) at 60 °C for 3 h (first, second, fourth, and fifth rounds) or for 16 h (third round), followed by centrifugation (20,400 × *g*, room temperature, 3 min). The supernatants were collected and 300 µL of CHCl_3_/CH_3_OH (1:2, v/v) was freshly added to the pellets each time. The supernatant of each fraction was dried and used as reversibly bound ceramide fractions. The pellets derived after the five-time elution were used as irreversibly bound fractions. The protein-bound ceramide fractions, reversibly bound fractions, and irreversibly bound fractions were each suspended in 300 µL of 1 M KOH in 95% CH_3_OH containing 2 µL of 1 µM *N*-(2’-(*R*)-hydroxypalmitoyl(*d*_9_)) D-*erythro*-sphingosine (Avanti Polar Lipids, Alabaster, AL, USA) as an internal standard and incubated at 60 °C for 1 h to release ω-OH ceramides from protein-bound ceramides. The samples were neutralized with 17.1 µL of 17.5 M acetic acid and mixed with 285 µL of CHCl_3_ and 239.4 µL of H_2_O successively. After centrifugation (20,400 × *g*, room temperature, 3 min), the lower, organic phase was collected, dried, dissolved in 500 µL CHCl_3_/CH_3_OH (1:2, v/v), and subjected to LC-MS/MS analysis (injection volume, 5 µL) as described below.

### Preparation of amino acid-conjugated protein-bound ceramides from mouse epidermis

Protein-bound ceramide fractions prepared from 10 mg of the epidermis as described above were suspended in 1 mL of CH_3_OH and transferred into a plastic tube. After centrifugation (20,400 × *g*, room temperature, 3 min), the pellets were washed with 500 µL of 50 mM Tris-HCl (pH 7.4), suspended in 135 µL of 50 mM Tris-HCl (pH 7.4) containing 1 mM DTT by sonication, treated with 15 µL of 10 mg/mL Pronase E [dissolved in 50 mM Tris-HCl (pH 7.4); Merck], and incubated at 37 °C for 2 h. The samples were then mixed with 562.5 µL CHCl_3_/CH_3_OH (1:2, v/v), 187.5 µL CHCl_3_, and 187.5 µL H_2_O, successively. After centrifugation (20,400 × *g*, room temperature, 3 min), the lower, organic phase was collected, dried, dissolved in 50 µL of CHCl_3_/CH_3_OH, and subjected to LC-MS/MS analysis (injection volume, 5 µL), as described below.

### Lipid analysis by LC-MS(/MS)

EE ceramides, EE-analog, and their Cys adducts were detected using an ultra-performance LC-coupled with tandem triple quadrupole mass spectrometer (Xevo TQ-S; Waters, Milford, MA, USA). Separation by LC was performed using a reverse phase column (Acquity UPLC CSH C18 column; length, 100 mm; particle size, 1.7 μm; inner diameter, 2.1 mm; Waters) and a binary gradient system comprising mobile phase A (CH_3_CN/H_2_O [3:2, v/v] containing 5 mM ammonium formate) and mobile phase B (CH_3_CN/2-propanol [1:9, v/v] containing 5 mM ammonium formate) at 55 °C with a flow rate of 0.3 mL/min. EE-analog and EE-analog–Cys conjugate were separated using the following gradient conditions: 0 min, 0% B; 0–3 min, linear gradient to 10% B; 3–10 min, linear gradient to 40% B; 10–18 min, linear gradient to 100% B; 18–23 min, 100% B; 23 min, return to 0% B; 23–25 min, 0% B. EE ceramides and EE-ceramide–Cys conjugates were separated using the following gradient conditions: 0 min, 40% B; 0–18 min, linear gradient to 100% B; 18–23 min, 100% B; 23 min, return to 40% B; and 23–25 min, 40% B. Ionization was performed by electrospray ionization and MS parameters were set as follows: capillary voltage, 2.5 kV; sampling cone, 30 V; source offset, 50 V; desolvation temperature, 650 °C; desolvation gas flow, 1,200 L/h; cone gas flow, 150 L/h.

EE-analog and EE-analog–Cys conjugate were detected using a single ion monitoring mode, in which molecular ions with respectively *m/z* = 235.5 and 356.6 were selected. Both were ionized by setting the cone voltage at 20 V. EE-ceramides, Cys-bound P-EO ceramides, and ω-OH ceramides were detected using MRM mode and/or precursor ion scanning mode. The *m/z* values of precursor ions and product ions and the voltage of collision energy in the MRM mode were set as per Table S1. In precursor ion scanning mode, the voltage of collision energy was set at 40 eV and the product ions with *m/z* = 264.3 and 309.0, corresponding to sphingosine and EE fatty acid moieties, respectively, were selected. Operation of the LC-MS(/MS) system and data analysis were performed using MassLynx software (version 4.1; Waters). Quantification of ω-OH ceramides was performed by calculating the ratio of the peak area for each ω-OH ceramide species to the internal standards.

### Statistical analysis

Data are presented as mean + SD. The significance of differences between the two groups was tested using the unpaired two-tailed Student’s *t*-test in Microsoft Excel (Microsoft, Redmond, WA, USA) and the significance of differences to controls in the presence of multiple groups was tested using Dunnett’s test in JMP13 (SAS Institute, Cary, NC, USA). A *P*-value of <0.05 was considered significant.

## Supplemental information

### Supplemental Figure legends

**Figure S1. Detection of EE-analog and its derivatives, related to Figure 3**

(A) EE-analog and its derivatives were separated by TLC and subjected to copper phosphate staining.

(B) The molecular ions with *m/z* = 235.5, 237.5, and 253.5 were detected via LC-MS using single ion monitoring mode.

**Figure S2. Detection of EE ceramides by LC-MS/MS, related to Figure 4**

EE ceramides were extracted from the protein-bound ceramide fractions and subjected to LC-MS/MS analysis using precursor ion scanning mode (scan range of *m/z*, 900–1,200) to detect precursor ions with the product ion of *m/z* = 264 (positive ion mode) (A, C) and those of *m/z* = 309 (negative ion mode). Total ion current chromatograms (A and B) and the mass spectra of the retention time of 12–14 min (B and C) are shown.

**Figure S3. Detection of conjugates of EE ceramide and amino acid, related to Figure 4**

EE ceramides were incubated with an amino acid (Cys, His, Arg, or Lys; 2 mM each) at 37 °C for 1 h and subjected to LC-MS/MS analysis using MRM mode. The proton adduct ions of each amino acid and EE ceramide containing C34:1 conjugates were selected as the precursor ions, and the product ion of *m/z* = 264 was selected.

**Figure S4. Absence of Glu-bound P-O ceramides in the epidermis, related to Figure 6**

Protein-bound ceramide fractions prepared from WT and *Cyp4f39* KO mouse epidermis (10 mg) were digested with pronase E (1 mg/mL) at 37 °C for 2 h and subjected to LC-MS/MS analysis using MRM mode. The proton adduct ions of Glu-bound P-O ceramide containing C32:1, C32:0, or C34:1 *N*-acyl moiety were selected as the precursor ions, and the product ion of *m/z* = 264 was selected.

**Figure S5. Scheme of EE-analog synthesis, related to Figure 2**

EE-analog was synthesized via the reaction scheme composed of six reaction steps as indicated. DMP, Dess-Martin periodinane; DBU, 1,8-diazabicyclo[5.4.0]undec-7-ene; DIBAL-H, diisobutylaluminium hydride; ^t^BuOOH, *tert*-Butyl hydroperoxide; Ti(O*^i^*Pr)_4_, tetraisopropyl orthotitanate; and MS4A, molecular sieves 4A.

