## Supplemental Figures and Table for "Determining the structure of protein-bound ceramides, essential lipids for skin barrier function"

### Figure S1

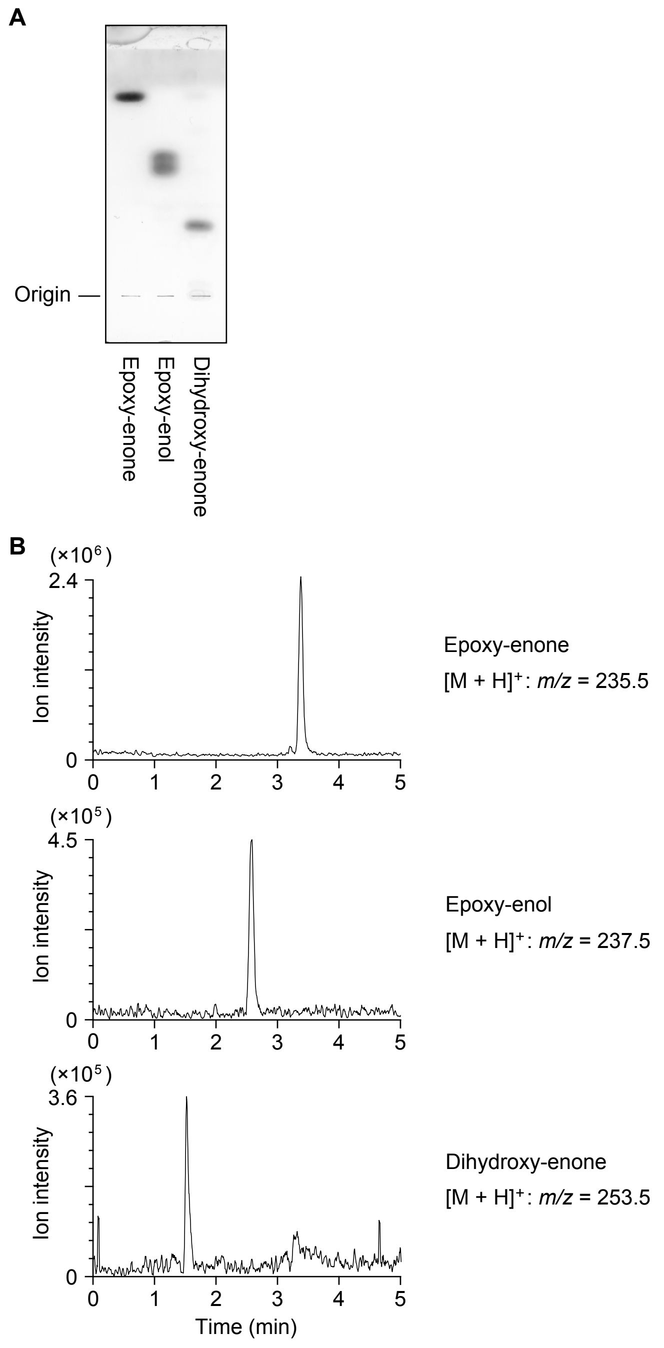
A
Origin
Epoxy-enone
Epoxy-enol
Dihydroxy-enone
B
(×10⁶)
2.4
Ion intensity
0
0 1 2 3 4 5
Epoxy-enone
[M + H]⁺: m/z = 235.5
(×10⁵)
4.5
Ion intensity
0
0 1 2 3 4 5
Epoxy-enol
[M + H]⁺: m/z = 237.5
(×10⁵)
3.6
Ion intensity
0
0 1 2 3 4 5
Time (min)
Dihydroxy-enone
[M + H]⁺: m/z = 253.5

### Figure S2

A

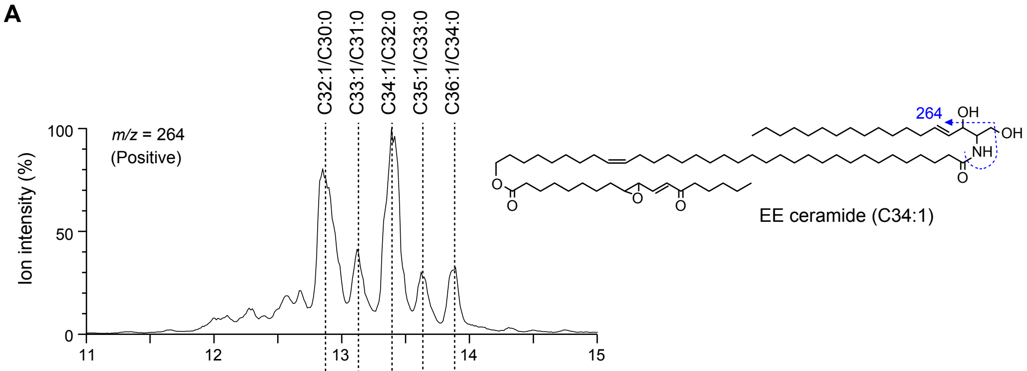


B

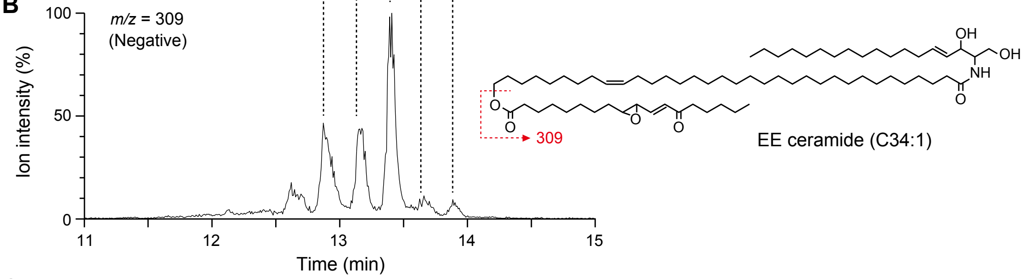


C

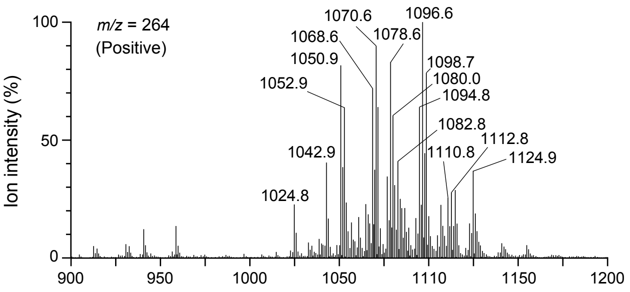


| $[M + H – H_2O]^+$ | $[M + H]^+$ | *N*-acyl |
|---|---|---|
| 1022.8 | 1040.8 | C30:1 |
| 1024.8 | 1042.8 | C30:0 |
| 1036.8 | 1054.8 | C31:1 |
| 1038.8 | 1056.8 | C31:0 |
| 1050.8 | 1068.8 | C32:1 |
| 1052.8 | 1070.8 | C32:0 |
| 1064.8 | 1082.8 | C33:1 |
| 1066.8 | 1084.8 | C33:0 |
| 1078.8 | 1096.8 | C34:1 |
| 1080.8 | 1098.8 | C34:0 |
| 1092.8 | 1110.8 | C35:1 |
| 1094.8 | 1112.8 | C35:0 |
| 1106.8 | 1124.8 | C36:1 |
| 1108.8 | 1126.8 | C36:0 |

D

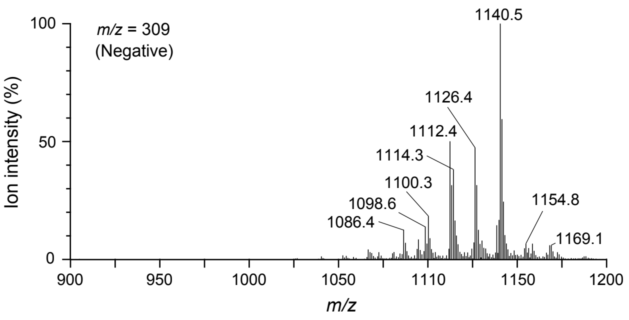


| $[M + HCOO]^–$ | *N*-acyl |
|---|---|
| 1084.8 | C30:1 |
| 1086.8 | C30:0 |
| 1098.8 | C31:1 |
| 1100.8 | C31:0 |
| 1112.8 | C32:1 |
| 1114.8 | C32:0 |
| 1126.8 | C33:1 |
| 1128.8 | C33:0 |
| 1140.8 | C34:1 |
| 1142.8 | C34:0 |
| 1154.8 | C35:1 |
| 1156.8 | C35:0 |
| 1168.8 | C36:1 |
| 1170.8 | C36:0 |

### Figure S3

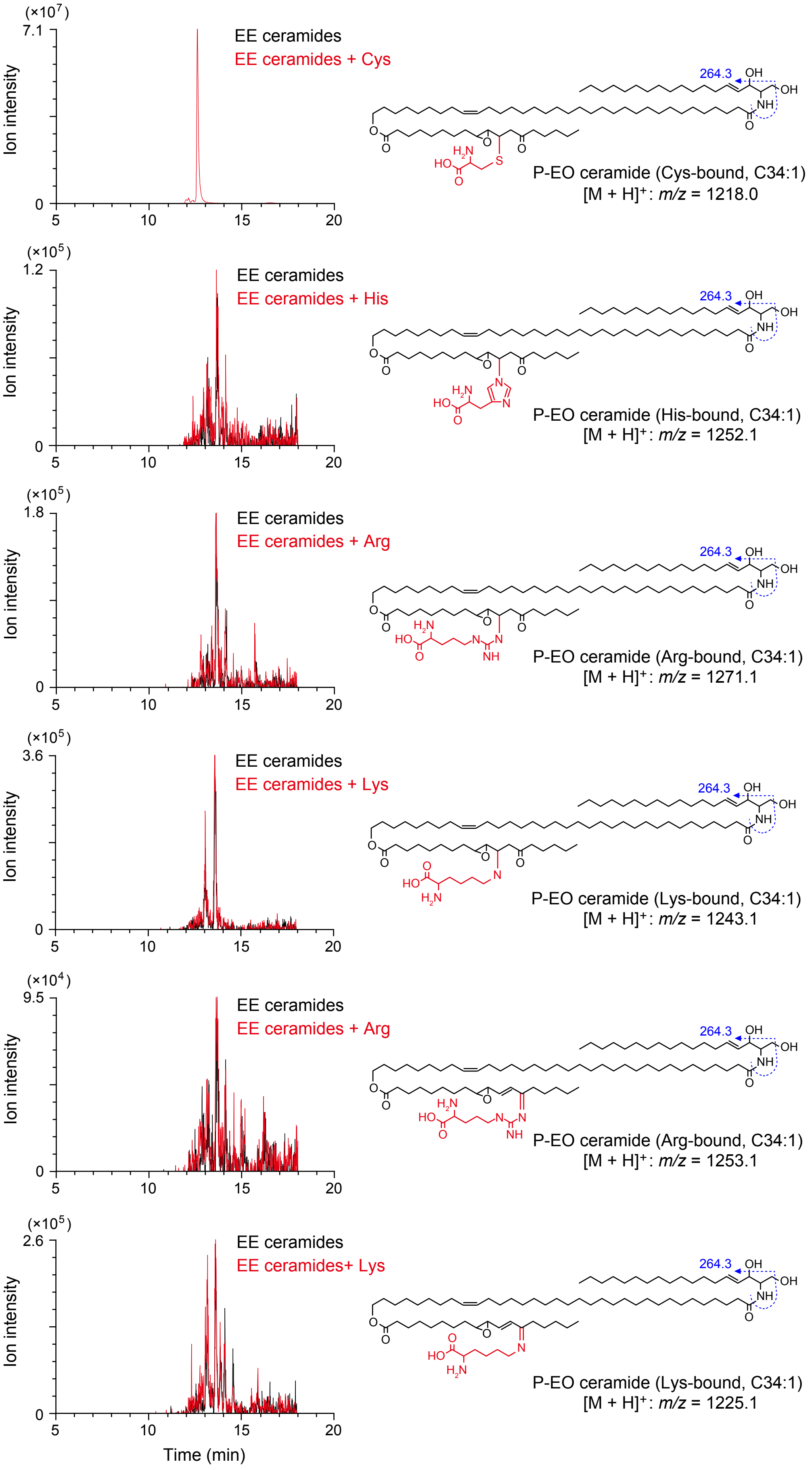
(×10^7)
7.1
Ion intensity
0
5
10
15
20
EE ceramides
EE ceramides + Cys
264.3
OH
OH
NH
O
O
O
O
O
H2N
HO
O
S
P-EO ceramide (Cys-bound, C34:1)
[M + H]+: m/z = 1218.0
(×10^5)
1.2
Ion intensity
EE ceramides
EE ceramides + His
P-EO ceramide (His-bound, C34:1)
[M + H]+: m/z = 1252.1
(×10^5)
1.8
Ion intensity
EE ceramides
EE ceramides + Arg
P-EO ceramide (Arg-bound, C34:1)
[M + H]+: m/z = 1271.1
(×10^5)
3.6
Ion intensity
EE ceramides
EE ceramides + Lys
P-EO ceramide (Lys-bound, C34:1)
[M + H]+: m/z = 1243.1
(×10^4)
9.5
Ion intensity
EE ceramides
EE ceramides + Arg
P-EO ceramide (Arg-bound, C34:1)
[M + H]+: m/z = 1253.1
(×10^5)
2.6
Ion intensity
EE ceramides
EE ceramides+ Lys
P-EO ceramide (Lys-bound, C34:1)
[M + H]+: m/z = 1225.1
Time (min)

### Figure S4

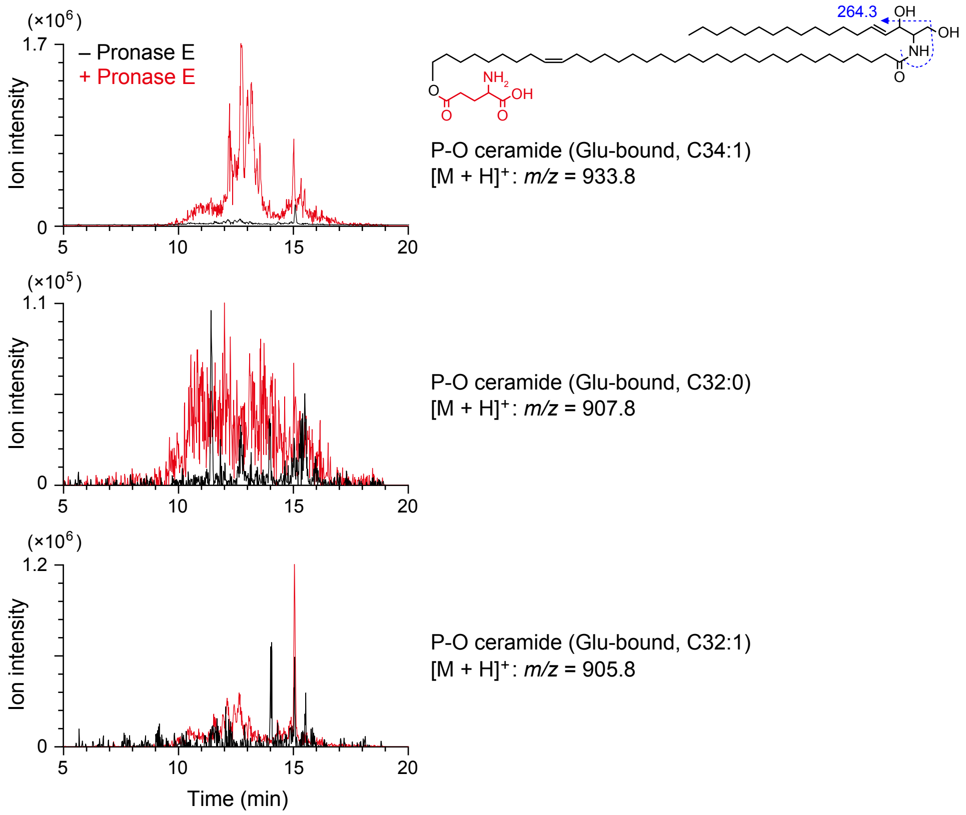
(×10^6)
1.7
0
– Pronase E
+ Pronase E
Ion intensity
5
10
15
20
264.3
OH
OH
NH
O
O
NH2
OH
O
O
P-O ceramide (Glu-bound, C34:1)
[M + H]+: m/z = 933.8
(×10^5)
1.1
0
P-O ceramide (Glu-bound, C32:0)
[M + H]+: m/z = 907.8
(×10^6)
1.2
0
P-O ceramide (Glu-bound, C32:1)
[M + H]+: m/z = 905.8
Time (min)

#### Figure S5

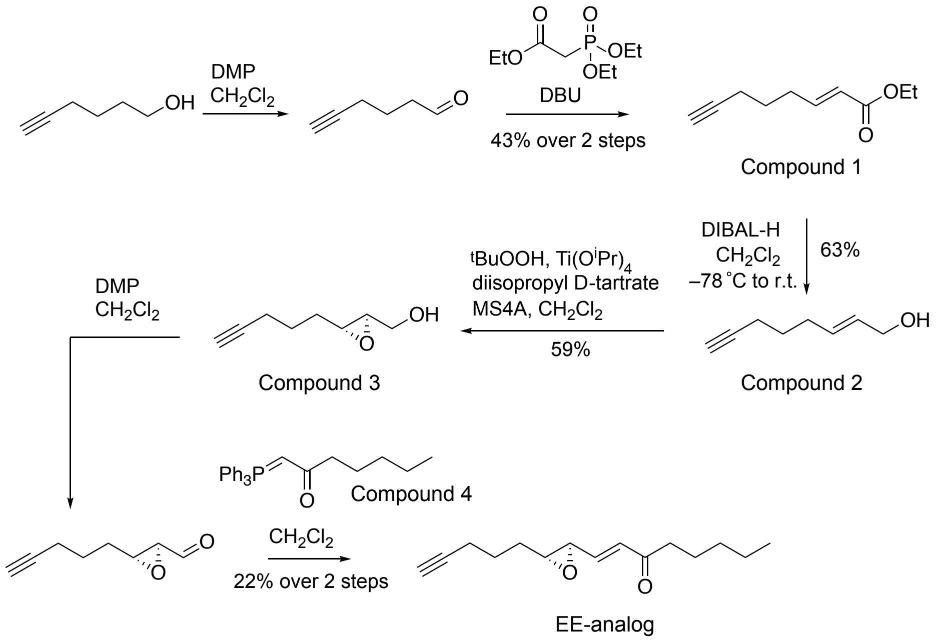

OH
DMP
$CH_2Cl_2$
O
EtO
P
OEt
OEt
DBU
43% over 2 steps
OEt
Compound 1
DIBAL-H
$CH_2Cl_2$
–78 ˚C to r.t.
63%
OH
Compound 2
$^t$BuOOH, Ti(O$^i$Pr)$_4$
diisopropyl D-tartrate
MS4A, $CH_2Cl_2$
59%
OH
Compound 3
DMP
$CH_2Cl_2$
$Ph_3P$
Compound 4
$CH_2Cl_2$
22% over 2 steps
EE-analog

#### Supplemental information

**Table S1.** MRM settings for detection of ceramide species in LC-MS/MS analyses

| Ceramides | *N*-acyl | Precursor ion (Q1) | | Product ion (Q3) | Cone voltage (V) | Collision energy (eV) |
|---|---|---|---|---|---|---|
| | | $[M + H - H_2O]^+$ | $[M + H]^+$ | | | |
| EE-ceramides | C30:0 | 1025.0 | 1043.0 | 264.3 | 30 | 35 |
| | C30:1 | 1023.0 | 1041.0 | 264.3 | 30 | 35 |
| | C32:0 | 1053.0 | 1071.0 | 264.3 | 30 | 40 |
| | C32:1 | 1051.0 | 1069.0 | 264.3 | 30 | 40 |
| | C34:0 | 1081.0 | 1099.0 | 264.3 | 30 | 40 |
| | C34:1 | 1079.0 | 1097.0 | 264.3 | 30 | 40 |
| | C36:0 | 1109.0 | 1127.0 | 264.3 | 30 | 45 |
| | C36:1 | 1107.0 | 1125.0 | 264.3 | 30 | 45 |
| EE-Cer–Cys conjugates | C30:1 | | 1164.0 | 264.3 | 30 | 35 |
| | C30:0 | | 1162.0 | 264.3 | 30 | 35 |
| | C32:1 | | 1192.0 | 264.3 | 30 | 40 |
| | C32:0 | | 1190.0 | 264.3 | 30 | 40 |
| | C34:1 | | 1220.0 | 264.3 | 30 | 40 |
| | C34:0 | | 1218.0 | 264.3 | 30 | 40 |
| | C36:1 | | 1248.0 | 264.3 | 30 | 45 |
| | C36:0 | | 1246.0 | 264.3 | 30 | 45 |
| ω-OH ceramides | C30:0 | 732.7 | 750.7 | 264.3 | 30 | 35 |
| | C30:1 | 730.7 | 748.7 | 264.3 | 30 | 35 |
| | C32:0 | 760.8 | 778.8 | 264.3 | 30 | 35 |
| | C32:1 | 758.8 | 776.8 | 264.3 | 30 | 35 |
| | C34:0 | 788.8 | 806.8 | 264.3 | 30 | 40 |
| | C34:1 | 786.8 | 804.8 | 264.3 | 30 | 40 |
| | C36:0 | 816.8 | 834.8 | 264.3 | 30 | 40 |
| | C36:1 | 814.8 | 832.8 | 264.3 | 30 | 40 |
| α-OH ceramides | $d_9$-C16:0 | 545.5 | 563.5 | 264.3 | 30 | 20 |
